# High Postoperative Monocyte Indicated Inferior Clinicopathological Characteristics and Worse Prognosis in Lung Adenocarcinoma or Squamous Cell Carcinoma after Lobectomy

**DOI:** 10.1101/395632

**Authors:** Yang Hai, Nan Chen, Wenwen Wu, Zihuai Wang, Feng Lin, Chenglin Guo, Chengwu Liu, Weiming Li, Lunxu Liu

## Abstract

**Background:** Peripheral monocyte count is an assessable parameter. Recently, evidence suggested an elevated preoperative monocyte counts predicting poor prognosis in malignancies. The aim of this study was to determine the prognostic effect of early postoperative blood monocyte count in patients with lung adenocarcinoma or squamous cell carcinoma following lobectomy.

**Methods:** We retrospectively reviewed patients with operated lung adenocarcinoma or squamous cell carcinoma from 2006 to 2011 in Western China Lung Cancer database. Univariate analysis on disease-free survival (DFS) and overall survival (OS) was performed using the Kaplan-Meier and log-rank tests, and multivariate analysis was conducted using the Cox proportional hazards regression model.

**Results:** There were 433 patients enrolled in our analysis. High postoperative elevated monocyte was associated with male gender (*P*<0.001), positive smoking history (*P*=0.005), and higher N stage (*P*=0.002) and higher tumor stage (*P*=0.026). Two-tailed log-rank test indicated patients with an early postoperative elevated monocyte count predicted a poor DFS and OS overall (*P*<0.001, *P*<0.001, respectively) as well as in subgroup analysis, and further presented as a promising independent prognostic factor for both DFS and OS (HR=2.991, 95%CI: 2.243-3.988, *P*<0.001; HR=2.705, 95%CI: 1.977-3.700, *P*<0.001, respectively) on multivariate analysis. However, no significance was detected for preoperative monocyte in multivariate analysis.

**Conclusions:** Elevated early postoperative peripheral monocyte count was an independent prognostic factor of poor prognosis and inferior clinicopathological features for patients with operable lung adenocarcinoma or squamous cell carcinoma by lobectomy.

## Introduction

Lung cancer was the leading cause of cancer-related death worldwide, with 5-year survival rates of less than 17% [1]. Among all types, non-small-cell lung cancer (NSCLC) accounts for approximately 85% of all lung cancers. Adenocarcinoma and squamous cell carcinoma account for approximately 40% and 25-30% of lung cancers, respectively [1,3]. In the aspect of treatment, if operable, surgery provided the best chance to cure NSCLC [2], which also called for a comprehensive perioperative evaluation system.

The hypothesis of the relationship between cancer and inflammation has triggered researcher interests though it was not new. Back to 1863, chronic inflammation was firstly regarded as the origin of tumor cells, and the response of the human body towards malignancies was proven to be closely related with inflammation [3]. Examples included tissue necrosis factor (TNF), interleukin (IL)-1, IL-6, matrix metalloproteinases, vascular endothelial growth factor, etc. As for peripheral blood cells, monocytes/macrophages, neutrophils, dendritic cells, and natural killer cells form the first line of immune defense in normal situations [4], and some of them have been regarded as “superstars” for oncologists. Specifically, in real clinical work, elevated preoperative monocyte counts have recently been shown to predict poor prognosis in various types of malignancies, including hepatic cell carcinoma, malignant lymphomas as well as lung adenocarcinoma [5,6,7]. However, it was still unclear if an elevated early postoperative monocyte count was associated with a poor prognosis in lung adenocarcinoma or squamous cell carcinoma after lobectomy. We focused on the early postoperative period because it was the most turbulent stage of the host immune system which may be caused by the surgical trauma and/or tumor removal effect, and we hypothesized that this change in immune environment related with tumor progression. The aim of this study was to evaluate whether an elevated early postoperative monocyte count predicted a poor prognosis in patients with operable lung adenocarcinoma or squamous cell carcinoma after lobectomy.

## Materials and Methods

### Study population

The data were retrieved from Western China Lung Cancer database (WCLC). We enrolled patients with operable lung adenocarcinoma or squamous cell carcinoma treated with lobectomy at West China Hospital, Sichuan University between 2006 and 2011. All patients were >18 years of age, with complete clinicopathological data, and proven to be lung adenocarcinoma or squamous cell carcinoma after surgery. Preoperative evaluation included physical examination, blood routine examination, tumor markers test, chest X-ray and computed tomography, brain magnetic resonance imaging (MRI), bone scintigraphy, and bronchoscopy and integrated positron emission tomography scan and CT (PET/CT) scan when necessary. The eligibility criteria were: 1) lobectomy with no microscopic residual tumor; 2) no preoperative chemotherapy and/or radiotherapy; 3) no previous history of other malignancies; 4) no evidence of infections such as pneumonia; 5) availability of laboratory data and follow-up information. The peripheral venous blood samples were collected from patients within one week before and within 4 days after surgery, the latter meant the first postoperative blood sample taken from the patient must be within 4 days, and only the first sample after surgery was recorded. Absolute peripheral blood count and the percentage were analyzed for each blood sample. Histological classification was made with reference to the latest WHO guideline [8]. The stages of lung cancer were confirmed based on the 7th edition of TNM classification of malignant tumors [9]. The study approval was granted by the Institutional Review Board at the West China Hospital, Sichuan University.

### Evaluation of clinicopathological factors

Baseline characteristics included age, sex, underlying diseases, smoking history, pathological stage, pathological tumor status, pathological lymph node status, and peripheral blood counts and the percentages. Blood samples used for analysis were taken within one week before and within 4 days after surgery performed within the clinical laboratory of our hospital via Cell-Dyn 3700 (Abbott Diagnostics, USA). Other data included the surgical date and procedures.

Treatment and follow-up

Lobectomy was performed on all patients with intent to cure, and systematic nodal dissection was carried out. The resection was done both macroscopically and histologically completely with a negative tumor margin and no evidence of distant metastasis. Patients were regularly followed up at outpatient department 1 month after surgery, every 3 months for the first year, every 6 months for the next four years, and once annually thereafter. Patients received a physical examination, blood routine examination, Chest and brain and upper abdomen CT scan at each follow-up. Bone scintigraphy was performed every 12 months. Particularly, all the patients treated in our department received phone call following-up regularly, during which we recorded their living status, tumor recurrence/metastasis condition, and sequential treatments such as chemotherapy and/or radiotherapy, etc. The patients were followed until August 31, 2017 or until they died.

### Statistical analysis

Receiver operating characteristic (ROC) curve was performed to search the best cut-off value for monocyte count to stratify patients at a high risk of tumor recurrence, distant metastasis, or death. In the ROC curve, the point with the maximum sensitivity and specificity was selected as the best cut-off value. Disease-free survival (DFS) was calculated from the date of surgery to the date of recurrence/metastasis or death with any cause, and overall survival (OS) was presented from the date of surgery to the date of death with any cause. DFS and OS were both censored at the date of the last follow-up. Fisher’s exact test or χ2-test for categorical variables and t-test for continuous variables were used to analyze the clinicopathological features for the two groups divided by the cut-off value of monocyte. Life table method was conducted to calculate 1, 3, 5-year survival rate. Survival curves were plotted using the Kaplan–Meier method and compared using the log-rank test. The prognostic factors of OS and DFS were analyzed by Cox proportional hazard model with univariate and multivariate analysis. Factors proven significant in the univariate analysis were included in the multivariate analysis. Subgroup analysis was used to further discriminate tumor prognosis between monocyte count and other prognostic factors: pathological stage, histological type, etc. All *P* values were two-tailed with less than 0.05 considered to be statistically significant. Statistical analysis was performed using SPSS (SPSS version 19.0, Chicago, IL, USA) and STATA 14.0 (STATA Corporation, College Station, TX, USA). Survival curves were drew by GraphPad Prism 5.0 (GraphPad Software, San Diego, CA).

## Results

The selection process of patients was showed in Figure 1. There were 1665 patients identified totally. 1232 patients were excluded for cancer type, pathological stage, or surgical procedure mismatch, previous history of malignancies or chemotherapy/radiotherapy, and those who combined with infections. Finally, there were 433 patients enrolled in our analysis. Among them, there were 278(64.2%) male and 155(35.8%) female. The average age of year’s old and standard difference were 60.6 and 10.0, respectively. Video-assisted thoracic surgery (VATS) was performed in 221(51.0%) patients compared with traditional thoracotomy surgery in 212(49.0%) patients. Specimens were histologically proven to be lung adenocarcinoma in 264(61.0%) patients and squamous cell carcinoma in 169(39.0%) patients. The details were presented in Table 1.

**Table 1.** Basic characteristics of clinicopathological features of patients with different monocyte counts

| Category | N (%) | Preoperative monocyte ($10^9/L$) | | Postoperative monocyte ($10^9/L$) | |
| --- | --- | --- | --- | --- | --- |
| | | <0.375 | $\geq 0.375$ | <0.845 | $\geq 0.845$ |
| Gender |  |  |  |  |  |
| Male | 278(64.2%) | 122 | 156 | 155 | 123 |
| Female | 155(35.8%) | 128 | 27 | 116 | 39 |
| Age | 433(100%) | 61.08±9.74 | 59.9±10.47 | 60.2±10.05 | 61.21±10.08 |
| Smoking history |  |  |  |  |  |
| Yes | 258(59.7%) | 115 | 143 | 148 | 110 |
| No | 174(40.3%) | 135 | 39 | 123 | 51 |
| Location |  |  |  |  |  |
| Left superior | 128(29.6%) | 74 | 54 | 80 | 48 |
| Left inferior | 62(14.3) | 39 | 23 | 40 | 22 |
| Right superior | 128(29.6%) | 79 | 49 | 80 | 48 |
| Right middle | 44(10.2) | 24 | 20 | 24 | 20 |
| Right inferior | 71(16.4%) | 34 | 37 | 47 | 24 |
| Procedure |  |  |  |  |  |
| VATS | 221(51.5%) | 139 | 82 | 148 | 73 |
| Thoracotomy | 212(49%) | 111 | 101 | 123 | 89 |
| T stage |  |  |  |  |  |
| T1 | 84(19.4%) | 58 | 26 | 50 | 34 |
| T2 | 279(64.4%) | 161 | 118 | 178 | 101 |
| T3 | 55(12.7%) | 26 | 29 | 31 | 24 |
| T4 | 15(3.5%) | 5 | 10 | 12 | 3 |
| N stage |  |  |  |  |  |
| N0 | 268(62.0%) | 158 | 110 | 184 | 84 |
| N1 | 71(16.4%) | 35 | 36 | 42 | 29 |
| N2 | 93(21.5%) | 56 | 37 | 45 | 48 |
| Stage |  |  |  |  |  |
| I | 205(47.3%) | 126 | 79 | 139 | 66 |
| II | 107(24.7%) | 59 | 48 | 68 | 39 |
| IIIA | 121(27.9%) | 65 | 56 | 64 | 57 |
| Histological type |  |  |  |  |  |
| Adenocarcinoma | 259(59.8%) | 181 | 78 | 168 | 91 |
| Squamous carcinoma | 174(40.2%) | 69 | 105 | 103 | 71 |
| Differentiation |  |  |  |  |  |
| Well | 11(2.6%) | 8 | 3 | 9 | 2 |
| Moderate | 261(61.7%) | 155 | 106 | 159 | 102 |
| Poor | 151(35.7%) | 80 | 71 | 98 | 53 |

T stage, tumour; N stage, node; VATS, Video-assistant thoracoscopic surgery; N, cases sample size
<sup>a</sup> Significance of Fisher's exact test, $\chi^2$ -test or t-test. $P < 0.05$ was considered statistically significant.

**Figure. 1.**
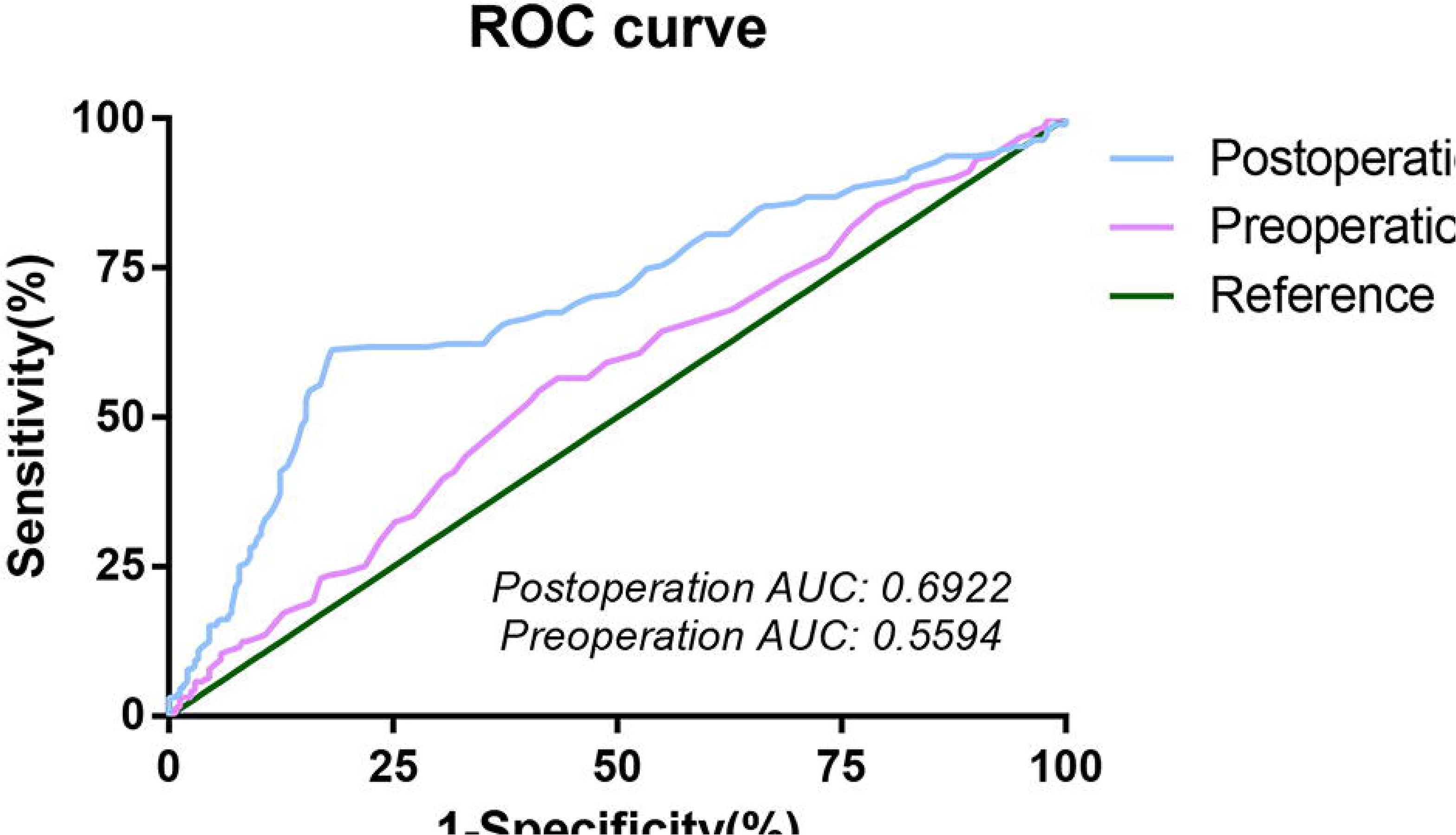
Flow chart of patient selection

By analyzing the ROC curve, the cut-off value for preoperative monocyte count was 0.375*10^9^/L with the area under the curve (AUC) of 0.566; while for postoperative monocyte count, the cut-off value was 0.845*10^9^/L with AUC of 0.692 (Figure 2). The baseline characteristics were stratified by high versus low preoperative monocyte and early postoperative monocyte count (Table 1). The preoperative monocyte was detected to be associated with male gender (*P*<0.001), smoking history (*P*<0.001), T stage (*P*=0.014) and histological type (*P*<0.001). For early postoperative group, compared to the low monocyte count group, male gender (*P*<0.001), positive smoking history (*P*=0.005), and higher N stage (*P*=0.002) and higher tumor stage (*P*=0.026) were more common found in high monocyte count group. The association between preoperative or postoperative monocyte and clinical characteristics were presented in Table 1 in details.

**Figure. 2.**
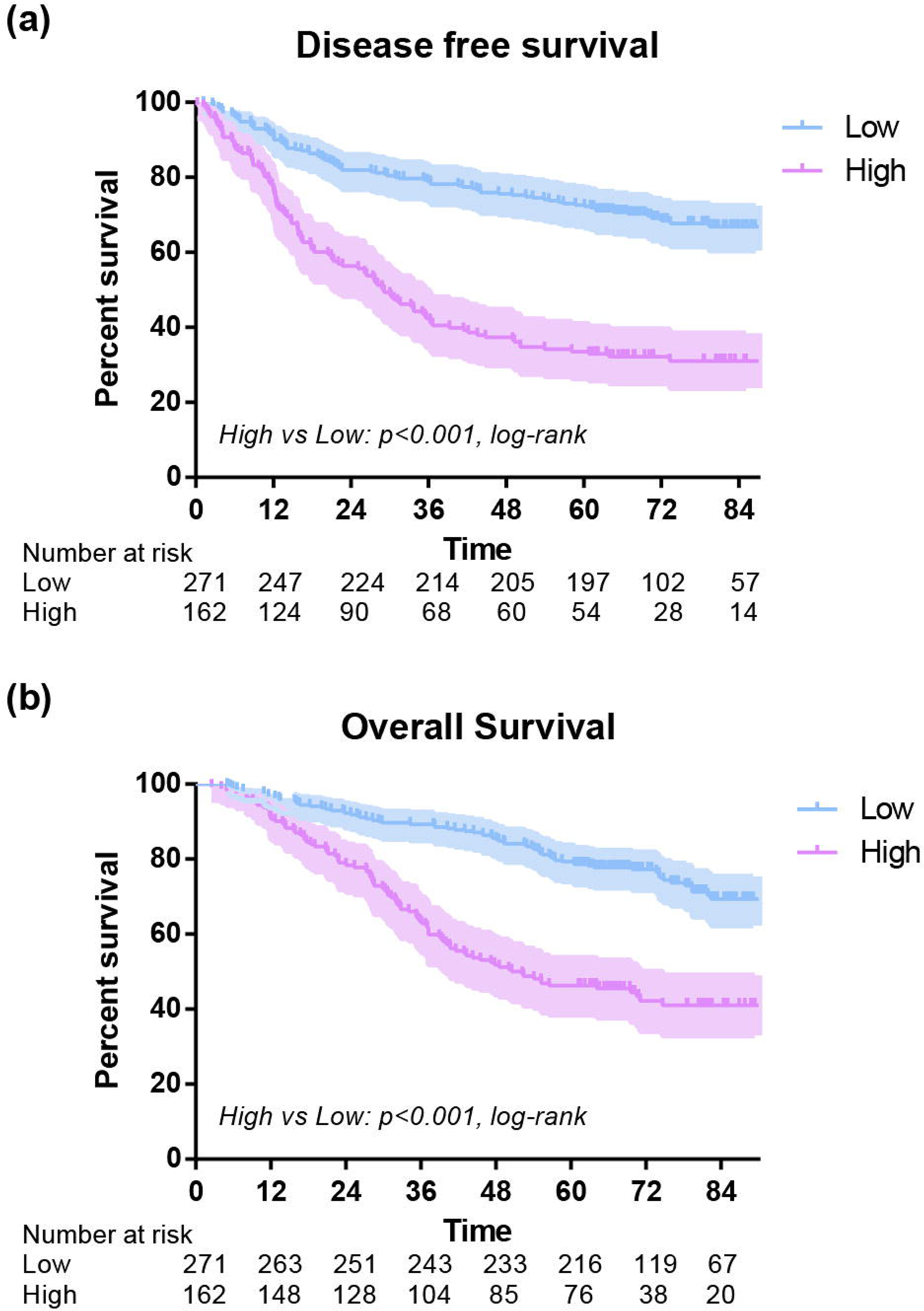
Receiver operating characteristic curve for determination of the cut-off value for monocyte. The cut-off value for preoperative monocyte count was 0.375*10^9^/L with area under the curve (AUC) of 0.566; while 0.845*10^9^/L with AUC of 0.692 for postoperative monocyte count.

We further performed Kaplan-Meier method to identify difference of survival rates between two groups stratified by monocyte count (low vs. high) of both preoperative and early postoperative group. For preoperative group, the 1-year OS of low and high subgroups were 95% and 93% (*P*=0.003), and the 1-year DFS of low and high subgroups were 82% and 85% (*P*=0.106). The 3-year OS of low and high subgroups were 82% and 74% (*P*=0.003), and the 3-year DFS of low and high subgroups were 67% and 60% (*P*=0.106), respectively. Additionally, the 5-year OS of low and high subgroups were 72% and 58% (*P*=0.003), and the 5-year DFS of low and high subgroups were 62% and 51% (*P*=0.106). As for the early postoperative group, the 1-year OS of low and high subgroups were 96% and 90% (*P*=0.004), and the 1-year DFS of low and high subgroups were 90% and 71% (*P*<0.001). The 3-year OS of low and high subgroups were 89% and 62% (*P*<0.001), and the 3-year DFS of low subgroup was 78% compared to 41% in high group (*P*<0.001). In addition, the 5-year OS of low and high subgroups were 79% and 46% (*P*<0.001), and the 5-year DFS of low and high subgroups were 72% and 33% (*P*<0.001). Overall, an early postoperative elevated monocyte count was significantly associated with poor OS (*P*<0.001) and DFS (*P*<0.001) (Figure 3). In further analysis, high level postoperative monocyte count was associated with poor OS and DFS in both adenocarcinoma and squamous carcinoma subgroups (Figure 4). These differences were also significant in subgroup analysis when stratified by gender, age, smoking history, stage, T stage, N stage and surgical procedure in postoperative group (Figure 5, Figure 6). On the contrary, no statistical significance was seen in subgroup analysis of preoperative group.

**Figure. 3.**
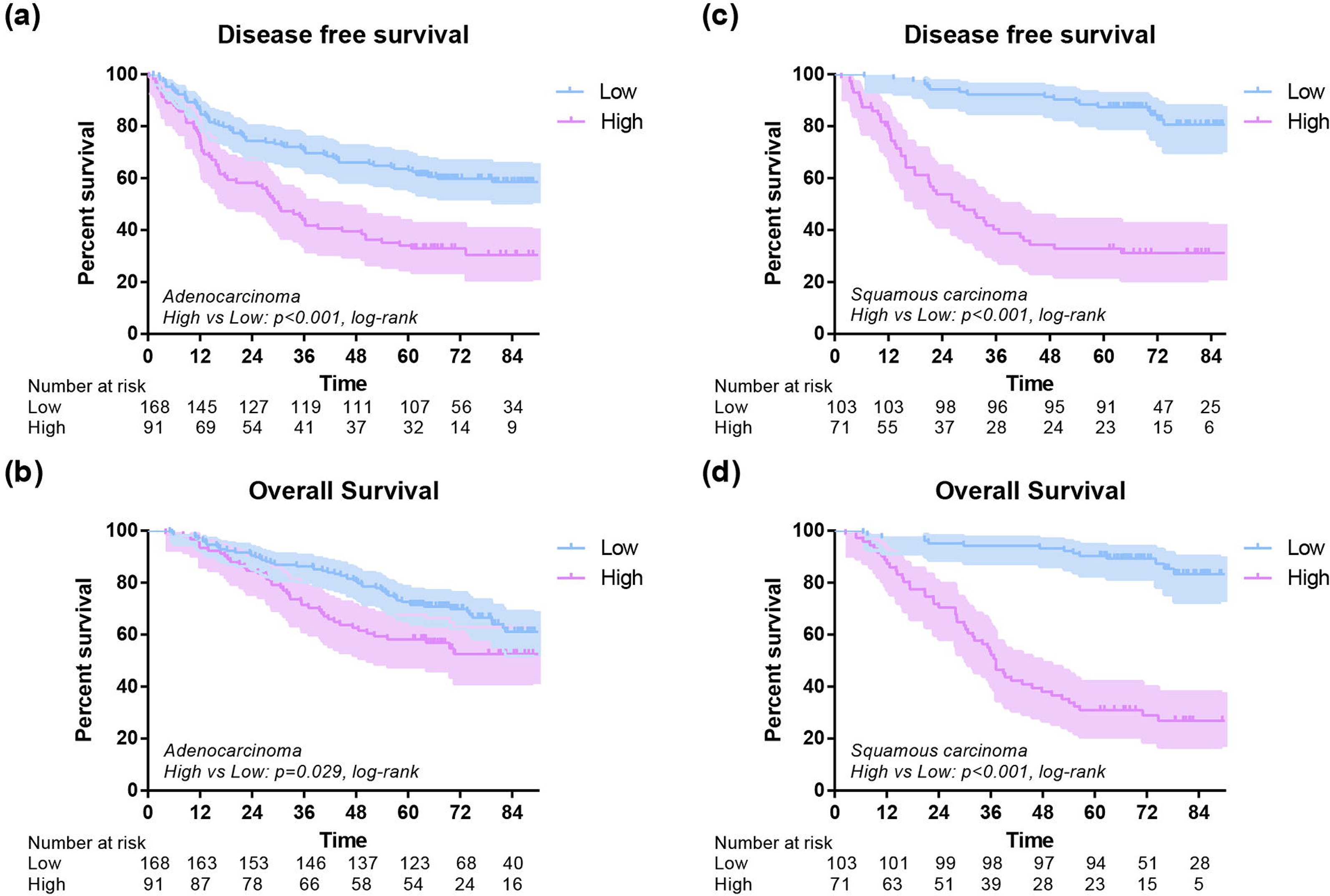
Disease-free survival(a) and overall survival(b) of high and low postoperative monocyte level.

**Figure. 4.**
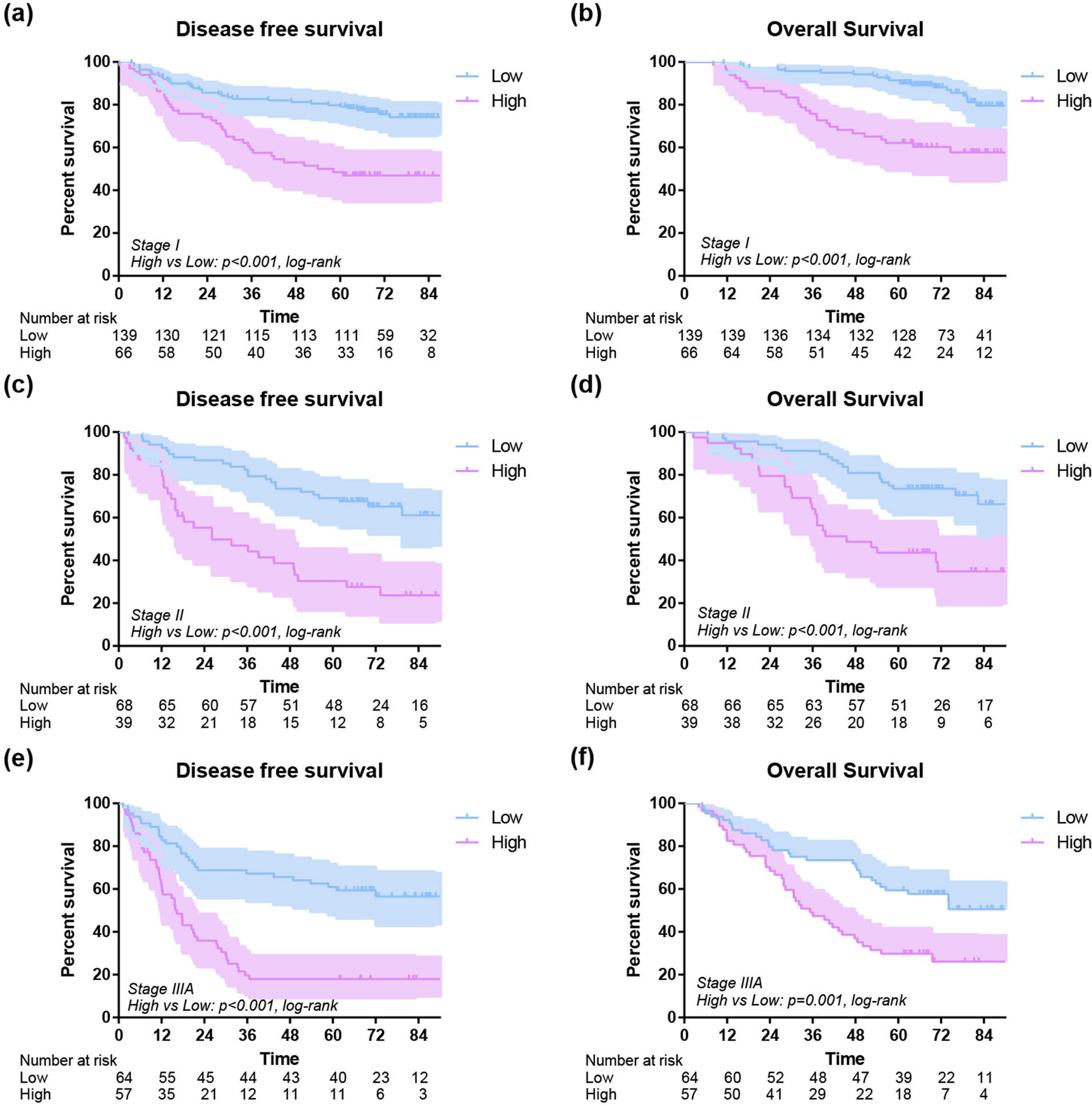
Disease-free survival and overall survival of 433 lung cancer patients with different monocyte stratified by histological type of adenocarcinoma(a,b) and squamous cell carcinoma(c,d).

**Figure. 5.**
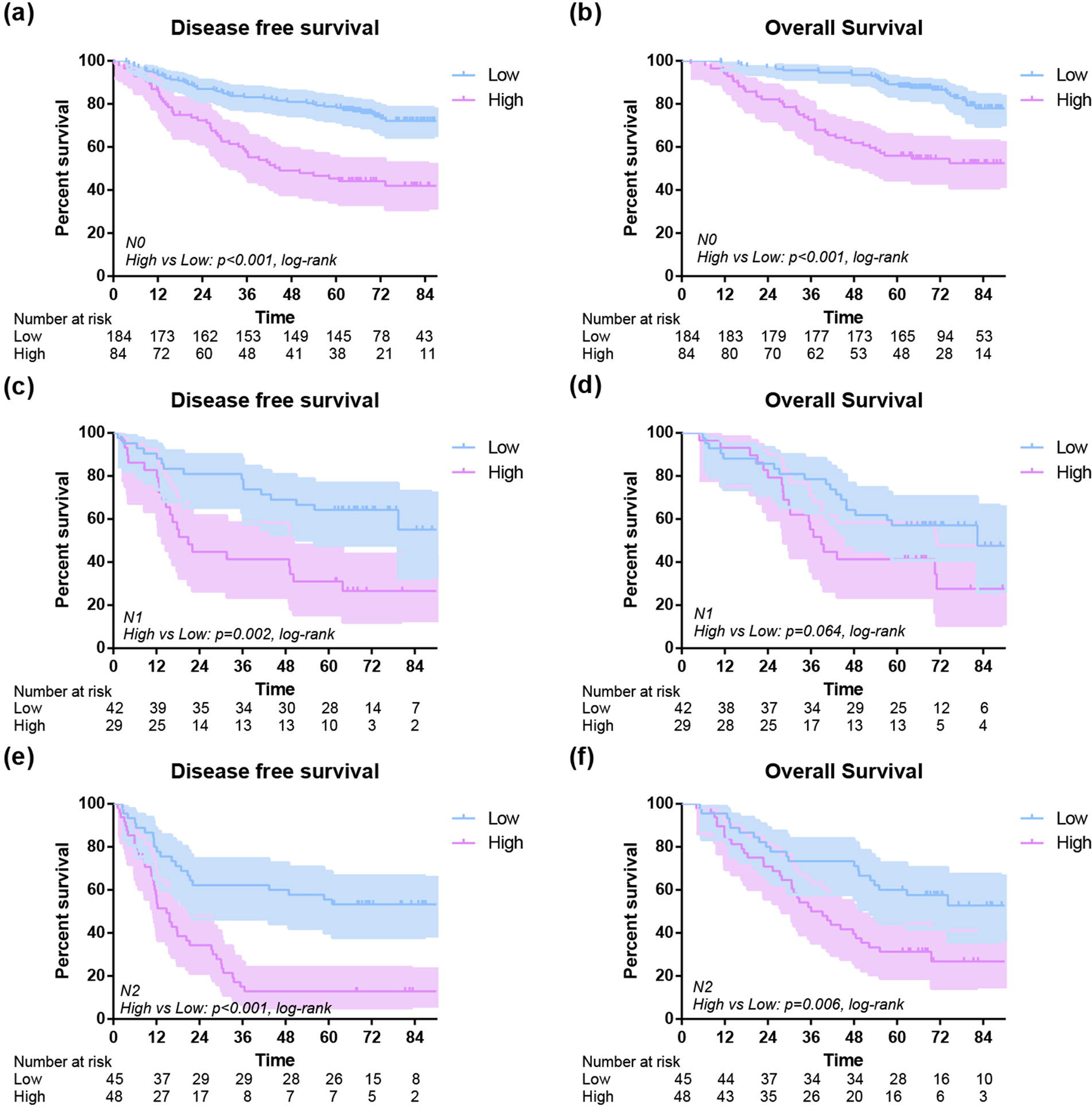
Disease-free survival and overall survival of 433 lung cancer patients with different monocyte stratified by tumor stage(stage I disease:a,b; stage II disease:c,d; stage IIIA disease:e,f.)

**Figure. 6.**
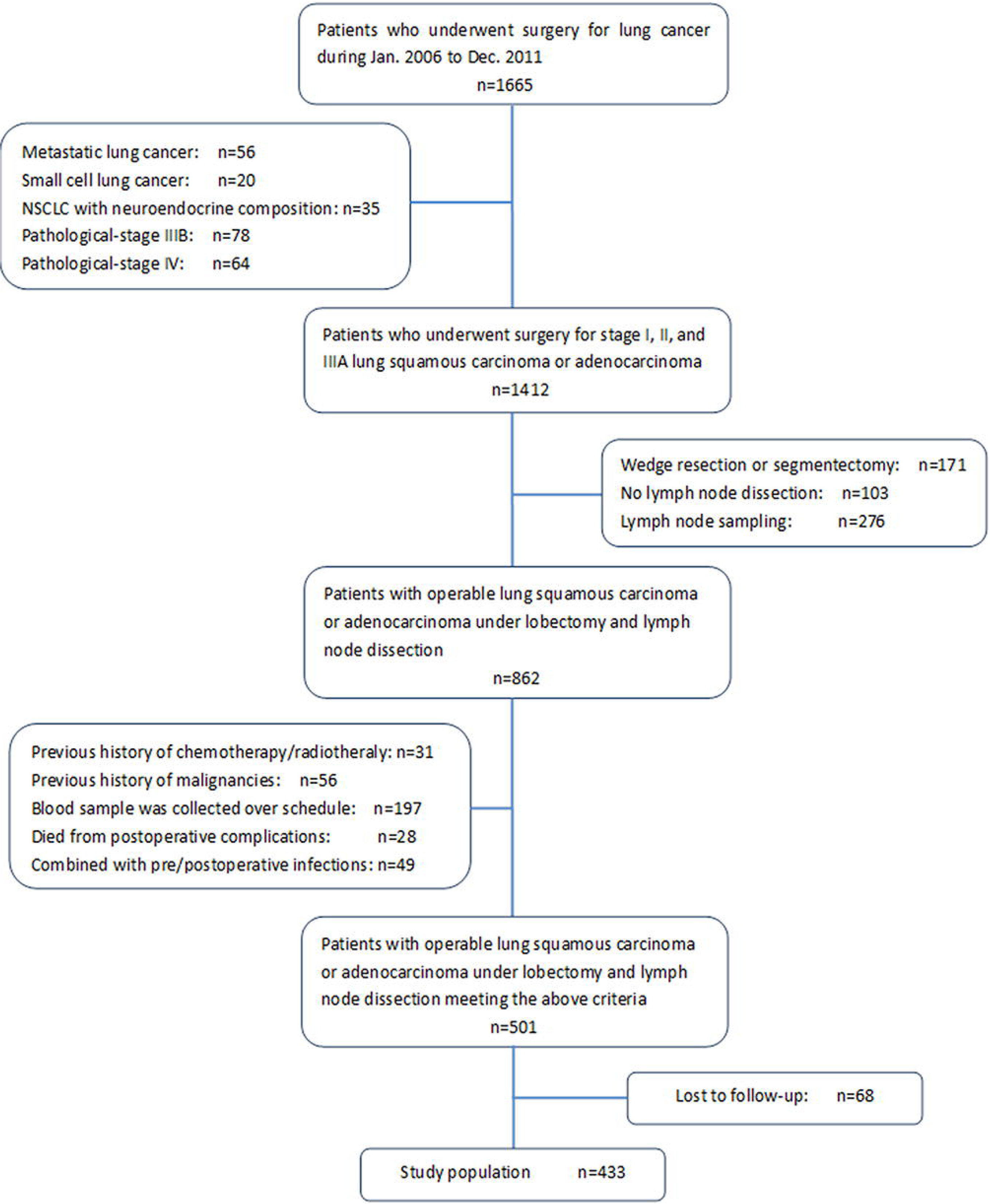
Disease-free survival and overall survival of 433 lung cancer patients with different monocyte stratified by N stage(N0 stage:a,b; N1 stage:c,d; N2 stage:e,f.)

Univariate prognostic analysis was performed to detect the prognostic significance of clinicopathological factors, and preoperative and early postoperative blood monocyte count (Table 2). N stage (*P*<0.001), tumor stage (*P*<0.001), histological type (*P*=0.014), preoperative (*P*=0.023) and postoperative monocyte count level (*P*<0.001) were significantly associated with DFS, while age (*P*=0.017), surgery procedure (*P*=0.002), T stage (*P*=0.003), N stage (*P*<0.001), tumor stage (*P*<0.001), preoperative(*P*=0.024) and postoperative monocyte count level (*P*<0.001) showed significant relationships with OS. The clinicopathological factors proved to be prognostic predictors in univariate analysis were included as covariates in further multivariate analysis (Table 3). Finally, thoracotomy (HR=1.520, 95%CI: 1.117-2.069, *P*=0.008), positive N status (HR=2.506, 95%CI: 1.625-3.864, P<0.001), squamous cell carcinoma (HR=0.633, 95%CI: 0.468-0.856, *P*=0.003) and high postoperative monocyte count (HR=2.991, 95%CI: 2.243-3.988, *P*<0.001) were risk factors with statistical significance for DFS. Correspondingly, age (HR=1.022, 95%CI: 1.005-1.038, *P*=0.009), thoracotomy (HR=1.700, 95%CI: 1.163-2.486, *P*=0.006), advanced tumor stage (HR=2.253, 95%CI: 1.178-4.309, *P*<0.001), and high postoperative monocyte count (HR=2.705, 95%CI: 1.977-3.700, *P*<0.001) were observed as risk factors for OS. However, preoperative monocyte was no more associated with either DFS or OS within multivariate analysis.

**Table 2.**
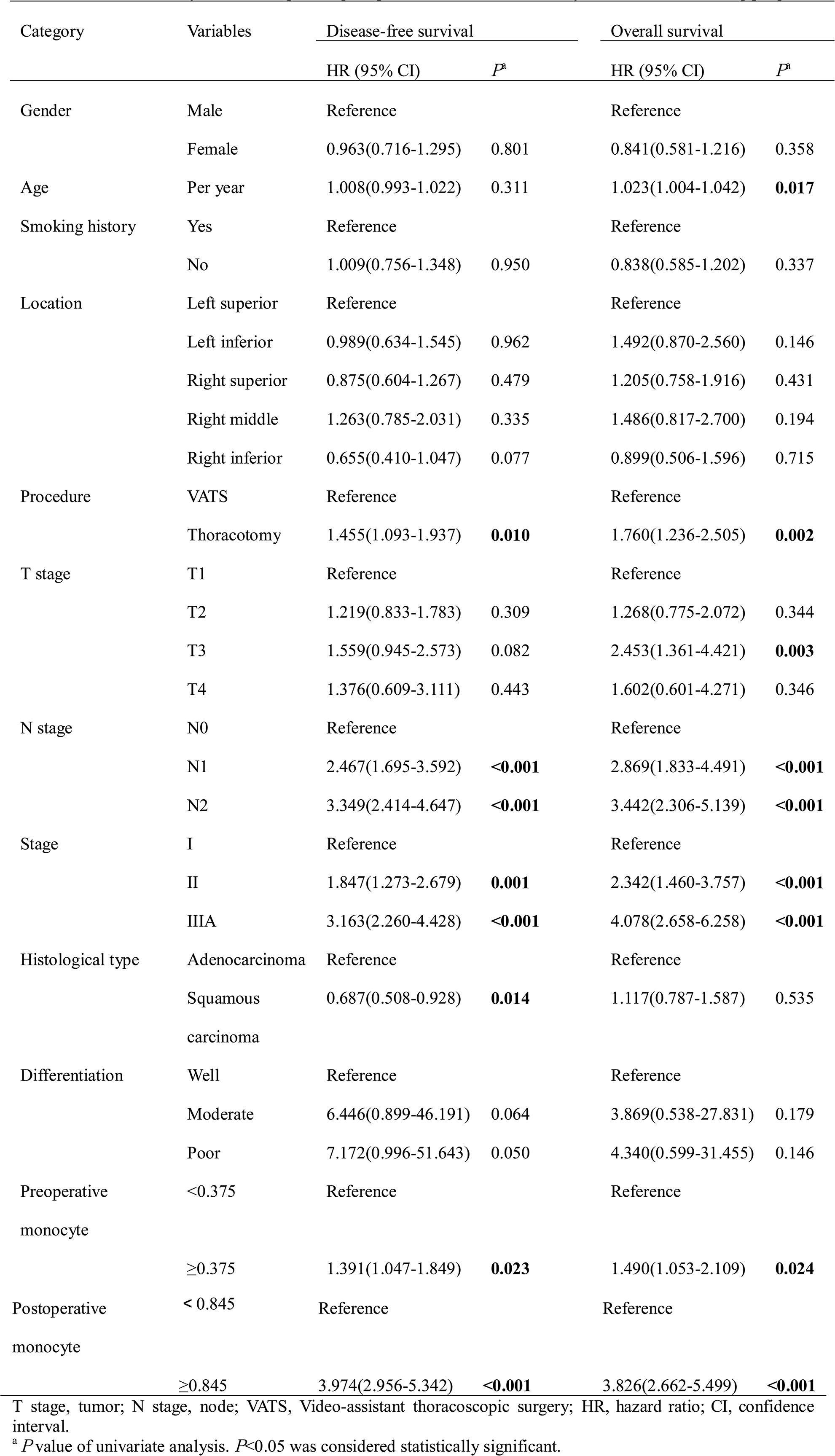
Univariate analysis of clinicopathological parameters and inflammatory biomarkers influencing prognosis

**Table 3.** Multivariate analysis of clinicopathological parameters and inflammatory biomarkers influencing prognosis

| Category | Variables | HR (95% CI) | *P*<sup>a</sup> |
| --- | --- | --- | --- |
| *Disease-free survival* |  |  |  |
| Procedure | VATS | Reference |  |
|  | Thoracotomy | 1.520(1.117-2.069) | **0.008** |
| N stage | N0 | Reference |  |
|  | N1 | 2.319(1.584-3.396) | **<0.001** |
|  | N2 | 2.584(1.845-3.620) | **<0.001** |
| Histological type | Adenocarcinoma | Reference |  |
|  | Squamous carcinoma | 0.561(0.405-0.779) | **0.001** |
| Postoperative monocyte | < 0.845 | Reference |  |
|  | ≥0.845 | 3.684(2.729-4.975) | **<0.001** |
| *Overall survival* |  |  |  |
| Age | Per year | 1.034(1.014-1.054) | **0.001** |
| Procedure | VATS | Reference |  |
|  | Thoracotomy | 1.700(1.163-2.486) | **0.006** |
| Stage | I | Reference |  |
|  | II | 2.228(1.383-3.592) | **0.001** |
|  | IIIA | 3.592(2.327-5.546) | **<0.001** |
| Postoperative monocyte | < 0.845 | Reference |  |
|  | ≥0.845 | 3.403(2.362-4.902) | **<0.001** |

N stage, node; VATS, Video-assistant thoroscopic surgery; HR, hazard ratio; CI, confidence interval.
<sup>a</sup> *P* value of multivariate analysis. *P*<0.05 was considered statistically significant.

## Discussion

Immune response was an essential component of tumor progression. Studying these responses within tumor microenvironment via several immune factors can further stratify the prognosis of cancer [10]. Back to 1997, Negus et al. had demonstrated a number of cells expressed chemokine receptors could infiltrate to tumor areas [11]. Later on, an increasing number of experimental studies found that inflammation played a crucial role in tumor progression. Inflammatory markers such as C-reactive protein in esophageal squamous cancer [12], Colony-stimulating factor-1 in mammary tumor [13], and inhibitors of metalloproteinases in NSCLC [14], all have been suggested as alternative markers for tumor progression [8]. Likewise, as easy assessable parameters, peripheral blood cell counts have also been regarded as predictors of tumor prognosis. An elevated neutrophil, monocyte and leukocyte counts were proven to be associated with poor survival in patients with metastatic melanoma [15].

Monocytes belonged to circulating peripheral blood cells that played the crucial role in immune response with the capability of differentiating into macrophages and antigen-presenting cells (APCs). Thus, they formed the first line of innate immune defense [16]. In addition, monocytes could activate T and B lymphocytes and further produce cytokines such as IL-12 and TNF-α to stimulate the immune response [17]. However, either overstimulation or immunosuppression of monocytes caused by surgical procedure and/or other external factors could disrupt the immune system [18,19]. Previous studies have found an elevated preoperative monocyte count demonstrated a poor prognostic factor in esophageal squamous cell carcinoma, mantle cell lymphoma, follicular lymphoma, and classical Hodgkin lymphoma, respectively [20,21,22,23]. Also, for the early postoperative period, Franke et al. discovered an increase in the absolute monocyte count but an impairment of monocyte function, indicating a decreased ability to synthesize IL-12 and TNF-α, to express HLA-DR, and to act as the APC [24]. To the best of our knowledge, this was the first study showing the prognostic significance of the early postoperative monocyte count in patients with lung adenocarcinoma or squamous cell carcinoma after lobectomy.

The mechanisms of elevated monocyte count related with poor prognosis of several kinds of tumors were still not clearly elucidated. There were several possible explanations. First, it was hypothesized that monocytes are attracted by several cytokines or chemokines to the tumor site and then differentiated into tumor-associated macrophages (TAMs), which further promoted those invading leukocytes to bring out potentials of angiogenesis, motility, and invasion [25,26]. Angiogenic signals from surrounding cells resulted in vasodilatation and increased vascular permeability [27,28], forming a vicious cycle for tumor progression. Second, human monocyte subsets were differentiated according to their surface CD14/CD16 expression as “majority/classical” (CD14++CD16-), “minority/non-classical” (CD14+CD16+) and the subset with pro-angiogenic feature (CD14++CD16+CCR2+) [29,30]. Among all subsets of the monocytes, the “majority/classical” accounts for approximately 90% of monocytes in healthy people [31]. However, Schauer et al. found that along with the increased number of monocytes, the major subsets shifted from CD16-to CD16+ after liver resection, showing a stronger potential of angiogenesis [32]. Correspondingly, while the classical monocytes were recruited to tumor sites, contributing to tumor macrophage content and promote tumor growth and metastasis, Richard N. Hanna et al. also found out the potential protective role of nonclassical “patrolling” monocytes tumor growth and metastasis [33]. This was also our next step to find out whether it will happen after lobectomy of lung cancer and what the exact type was and the alternatives of the function in immunity. Third, peripheral monocytes grow into TAMs when entering tumor areas. TAMs are classified into two phenotypes: M1 and M2. Activated M1 macrophages have the anti-tumor response, while M2 macrophages, activated by tumor-derived cytokines, were suitable for tumor development [34,35]. A previous study has reported that circulating macrophages predict tumor recurrence after surgery in patients with NSCLC [36], while in this study, an elevated early postoperative peripheral monocyte count was significantly associated with a poor prognosis in patients with lung adenocarcinoma or squamous cell carcinoma after lobectomy. These results might suggest a complex association between peripheral TAMs (M2 type) and monocytes, which still called further studies to verify. Moreover, the reason why we focused on the early postoperative period--the most turbulent stage of host immune system caused by surgical trauma and/or tumor removal effect probably was that we hypothesized that change in this period of immune environment might relate with tumor progression more closely than that of the preoperative data.

In the present study, an elevated early postoperative monocyte count was shown to predict a poor DFS and OS both in the univariate and multivariate analysis. In addition, as shown in the subgroup analysis, the monocyte count was found to be significantly associated with poor prognosis when stratified by gender, age, smoking history, TNM stage, surgical procedure and histological type. All of the above indicated that an elevated early postoperative blood monocyte count to be a very strong prognostic factor in tumor progression.

Limitations of the current study are inherent to its design, including the retrospective data collection and several confounding factors when comparing postoperatively. Moreover, the small number of patients, especially the cases with endpoints, also limited the conclusion of the current study.

## Conclusion

The present study supported the prognostic significance of early postoperative peripheral blood monocyte count in patients with operable lung adenocarcinoma or squamous cell carcinoma after lobectomy in both OS and DFS. This easily measured blood parameter may provide useful information for the clinicians to stratify patients. Further investigations were still needed to figure out the oncological significance of monocyte and its subsets, the association with host inflammatory microenvironment.

## Abbreviations

OS: overall survival
DFS: disease-free survival
NSCLC: non-small cell lung cancer
TNF: tissue necrosis factor
IL: interleukin
MRI: magnetic resonance imaging
PET-CT: positron emission tomography scan and CT
ROC: receiver operating characteristic
VATS: video-assisted thoracic surgery
AUC: area under the curve
APCs: antigen-presenting cells
TAMs: tumor-associated macrophages

## Declarations

### Ethics approval and consent to participate

The study approval was approved by the Institutional Review Board at the West China Hospital, Sichuan University. All participants have signed the consent to patients for participating in this study.

### Consent for publication

Not applicable

### Availability of data and material

The datasets generated and analysed during the current study are not publicly available due to confidential agreement but are available from the corresponding author on reasonable request.

### Authors’ contributions

LXL conceptualized the study and critically read the manuscript. YH, NC, WWW, ZHW, FL, CLG and CWL performed and/or assisted surgery, managed patients, and participated in data analysis. YH and NC wrote the manuscript. All authors read and approved the final manuscript

## Acknowledgements

Not applicable

## References

1. Siegel RL, Miller KD, Jemal A. Cancer statistics. CA Cancer J Clin. 2018 Jan; 68(1):7–30.

2. Non-Small Cell Lung Cancer. In: Detailed Guide. The American Cancer Society. 2016. http://www.cancer.org/cancer/lungcancer-non-smallcell/detailedguide/index. Accessed 25 October 2016

3. Balkwill F, Mantovani A. Inflammation and cancer: back to Virchow. Lancet. 2001;357: 539–545.

4. Coussens LM, Zitvogel L, Palucka AK. Neutralizing tumor-promoting chronic inflammation: a magic bullet. Science. 2013;339:286–91.

5. Shen SL, Fu SJ, Huang XQ, Chen B, Kuang M, Li SQ, Hua YP, Liang LJ and Peng BJ. Elevated preoperative peripheral blood monocyte count predicts poor prognosis for hepatocellular carcinoma after curative resection. BMC Cancer. 2014;14:744.

6. Porrata LF, Ristow K, Habermann TM, Witzig TE, Colgan JP, Inwards DJ, Ansell SM, Micallef IN, Johnston PB, Nowakowski GS, Thompson C, Markovic SN. Peripheral blood lymphocyte/monocyte ratio at diagnosis and survival in classical Hodgkin’s lymphoma. Haematologica. 2012;97:262–9.

7. Kumagai S, Marumo S, Shoji T, Sakuramoto M, Hirai T, Nishimura T, Arima N, Fukui M, Huang CL. Prognostic impact of preoperative monocyte counts in patients with resected lung adenocarcinoma. Lung Cancer. 2014;85:457–464.

8. Travis WD, Bramvilla E, Muller-Hermelink Hk, Harris CC. Tumours of the lung, pleura, thymus and heart. 4th ed. Lyon: International Agency for Research on Cancer; 2015

9. Sobin LH, Gospodarowicz MK, Wittekind C. International Union Against Cancer (UICC): TNM classification of malignant tumours. 7th ed. UK: John Willkey and Sons Ltd; 2010

10. Dvorak HF, Weaver VM, Tlsty TD, Bergers G. Tumor microenvironment and progression. J Surg Oncol. 2011;103:468–474.

11. Negus RPK, Stamp GWQ, Hadley J, Balkwill FR. Quantitative assessment of the leucocyte infiltrate in ovarian cancer and its relationship to the expression of C-C chemokines. Am J Pathol. 1997;150:1723–34.

12. Badakhshi H, Kaul D, Zhao KL. Association between the inflammatory biomarker, C-reactive protein, and the response to radiochemotherapy in patients with esophageal cancer. Mol Clin Oncol. 2016;4:643–647.

13. Domínguez SA Sierra FE, Puig KA, Pérez MB, Gómez AF, Corcuera MT, Sánchez MP, Corbí AL. Dendritic cell-specific ICAM-3-grabbing nonintegrin expression on M2-polarized and tumorassociated macrophages is macrophage-CSF dependent and enhanced by tumor-derived IL-6 and IL-10. J Immunol. 2011;186:2192–200.

14. Gouyer V, Conti M, Devos P, Zerimech F, Copin MC, Créme E, Wurtz A, Porte H, Huet G. Tissue inhibitor of metalloproteinase 1 is an independent predictor of prognosis in patients with nonsmall cell lung carcinoma who undergo resection with curative intent. Cancer. 2005; 103:1676–1684.

15. Gandini S, Ferrucci PF, Botteri E, Tosti G, Barberis M, Pala L, Battaglia A, Clerici A, Spadola G, Cocorocchio E, Martinoli C. Prognostic significance of hematological profiles in melanoma patients. Int J Cancer. 2016;139:1618–25.

16. Kyoizumi S, Kubo Y, Kajimura J, Yoshida K, Hayashi T, Nakachi K, Young LF, Moore MA, van den Brink MR, Kusunoki Y. Linkage between dendritic and T cell commitments in human circulating hematopoietic progenitors. Immunol. 2014;192:5749–60

17. Chen HW, Chen HY, Wang LT, Wang FH, Fang LW, Lai HY, Chen HH, Lu J, Hung MS, Cheng Y, Chen MY, Liu SJ, Chong P, Lee OK, Hsu SC. Mesenchymal stem cells tune the development of monocyte-derived dendritic cells toward a myeloid-derived suppressive phenotype through growth-regulated oncogene chemokines. J Immunol. 2013;190:5065 –77.

18. Nguyen BA, Fiorentino F, Reeves BC, Baig K, Athanasiou T, Anderson JR, Haskard DO, Angelini GD, Evans PC. Mini Bypass and Proinflammatory Leukocyte Activation: A Randomized Controlled Trial. Ann Thorac Surg. 2016;101:1454–63.

19. Fragiadakis GK, Gaudillière B, Ganio EA, Aghaeepour N, Tingle M, Nolan GP, Angst MS. Patient-specific Immune States before Surgery Are Strong Correlates of Surgical Recovery. Anesthesiology. 2015;123:1241–55.

20. Kumagai S, Marumo S1, Shoji T, Sakuramoto M, Hirai T, Nishimura T, Arima N, Fukui M, Huang CL. Prognostic impact of preoperative monocyte counts in patients with resected lung adenocarcinoma. Lung Cancer. 2014;85:457–464.

21. von Hohenstaufen KA, Conconi A, de Campos CP, Zucca E. Prognostic impact of monocyte count at presentation in mantle cell lymphoma. Br J Haematol. 2013;162:465–73.

22. Wilcox RA, Ristow K, Habermann TM, Inwards DJ, Micallef IN, Johnston PB, Colgan JP, Nowakowski GS, Ansell SM, Witzig TE, Markovic SN, Porrata L. The absolute peripheral monocyte count is associated with overall survival in patients newly diagnosed with follicular lymphoma. Leuk Lymphoma. 2012;53:575–80.

23. Porrata LF, Ristow K, Colgan JP, Habermann TM, Witzig TE, Inwards DJ, Ansell SM, Micallef IN, Johnston PB, Nowakowski GS, Thompson C, Markovic SN. Peripheral blood lymphocyte/monocyte ratio at diagnosis and survival in classical Hodgkin’s lymphoma. Haematologica. 2012; 97:262–9.

24. Franke A, Lante W, Fackeldey V, Becker HP, Kurig E, Zöller LG, Weinhold C, Markewitz A. Pro-inflammatory cytokines after different kinds of cardio-thoracic surgical procedures: is what we see what we know. Eur J Cardiothorac Surg. 2015;28:569–575.

25. Wu H, Xu JB, He YL, Peng JJ, Zhang XH, Chen CQ, Li W, Cai SR. Tumor-associated macrophages promote angiogenesis and lymphangiogenesis of gastric cancer. J Surg Oncol. 2012;106:462–468.

26. Jaipersad AS, Lip GY, Silverman S, Shantsila E. The role of monocytes in angiogenesis and atherosclerosis. J Am Coll Cardiol. 2014;63: 1–11.

27. Kimura H, Esumi H. Reciprocal regulation between nitric oxide and vascular endothelial growth factor in angiogenesis. Acta Biochim Pol. 2003;50:49–59.

28. Pamukcu B, Lip GY, Devitt A, Griffiths H, Shantsila E. The role of monocytes in atherosclerotic coronary artery disease. Ann Med. 2010;42:394–403.

29. González D, Domínguez S, Nieto C. Atypical Activin A and IL-10 Production Impairs Human CD16+ Monocyte Differentiation into Anti-Inflammatory Macrophages. J Immunol. 2016;196:1327–37.

30. Shantsila E, Wrigley B, Tapp L, Apostolakis S, Montoro-Garcia S, Drayson MT, Lip GY. Immunophenotypic characterization of human monocyte subsets: possible implications for cardiovascular disease pathophysiology. J Thromb Haemost. 2011;9:1056–66.

31. Sbrana S, Parri MS, De Filippis R, Gianetti J, Clerico A. Monitoring of monocyte functional state after extracorporeal circulation: A flow cytometry study. Cytometry. 2004;58B:17.

32. Schauer D, Starlinger P, Zajc P. Monocytes with angiogenic potential are selectively induced by liver resection and accumulate near the site of liver regeneration. BMC Immunol. 2014;15: 50.

33. Hanna RN, Cekic C, Sag D, Tacke R, Thomas GD, Nowyhed H, Herrley E, Rasquinha N, McArdle S, Wu R, Peluso E, Metzger D, Ichinose H, Shaked I, Chodaczek G, Biswas SK, Hedrick CC. Patrolling monocytes control tumor metastasis to the lung.Science. 2015; 33. 350:985–90.

34. Mantovani A, Sozzani S, Locati M, Allavena P, Sica A. Macrophage polarization: tumor-associated macrophages as a paradigm for polarized M2 mononuclear phagocytes. Trends Immunol. 2002;23:549–55.

35. Qian BZ, Pollard JW. Macrophage diversity enhances tumor progression andmetastasis. Cell. 2010;141:39–51.

36. Maeda R, Ishii G, Neri S, Aoyagi K, Haga H, Sasaki H, Nagai K, Ochiai A. Circulating CD14+CD204+ cells predict postoperative recurrence in non-small-cell lung cancer patients. J Thorac Oncol. 2014;9:179–8.

